# Exploring human papilloma virus-induced cytological and molecular alterations: a hybrid study on inflammatory pathways and therapeutic strategies

**DOI:** 10.1101/2025.09.05.674423

**Authors:** Mahya Nasrollahi, Mojgan Bandehpour, Hojun Kim

## Abstract

Cervical cancer remains a persistent and critical challenge in global health, primarily driven by chronic infections with high-risk human papillomavirus (HPV) types. This virus strategically exploits inflammatory signaling pathways, such as NF-κB, and modulates key enzymes like COX-2, promoting chronic inflammation and cellular alterations that significantly elevate the risk of carcinogenesis. A precise and comprehensive understanding of these pathways, combined with insights from advanced studies, is imperative for developing effective strategies to manage inflammation and mitigate the adverse impacts of HPV on cellular health. This study follows our previous research, which established HPV genotyping and related analyses, providing a foundation for the current investigation. It confirms a strong association between high-risk HPV genotypes and significant cytological and inflammatory alterations. Among individuals infected with high-risk HPV (16, 18, 52), 90 % exhibited notable cellular changes and severe inflammatory responses, compared to 45.5% in low-risk HPV cases and 59.3% in samples containing other HPV genotypes. In the 20 -30-year age group, 100% of samples displayed endocervical/metaplastic cells, with 70% exhibiting substantial cytological abnormalities, highlighting an increased susceptibility of younger individuals to HPV-induced epithelial disruptions. These findings emphasize the critical role of NF-κB and AhR pathways in modulating cellular responses, providing a foundation for targeted therapeutic interventions. Regulating NF-κB activity and microbiota-driven AhR modulation emerge as promising therapeutic strategies for mitigating inflammation and cellular damage associated with HPV infections. By integrating cytological observations with molecular analyses, this study presents a comprehensive perspective on HPV-induced pathophysiology, reinforcing the necessity for precision-based therapeutic approaches to effectively manage HPV-driven cellular and inflammatory alterations.

**Author Summary:** In this study, we investigated how human papillomavirus (HPV) triggers inflammation and cellular changes that can lead to cervical cancer. We combined laboratory analysis of patient samples with a review of existing research to map out the molecular pathways involved.

We found that infections with high-risk HPV types (like 16, 18, and 52) cause severe inflammation and cellular damage, especially in younger women aged 20-30. Specifically, 90% of samples with high-risk HPV showed significant inflammatory changes, compared to only 45.5% in low-risk cases.

Our results highlight the critical role of two key pathways—NF-κB and AhR—in driving HPV-induced inflammation. Targeting these pathways with anti-inflammatory compounds (like protopine) or immune-modulating agents (like immunobiotics) could offer new strategies to prevent or treat HPV-related cervical abnormalities.

This work underscores the importance of early HPV screening and personalized therapeutic interventions to reduce the risk of cervical cancer, particularly in high-risk populations.

## Introduction

Human papillomavirus (HPV) is a significant public health concern due to its established role as a causative agent in the development of cervical cancer and other anogenital malignancies. HPVs are a diverse group of more than 200 related DNA viruses, many of which have been classified as high-risk due to their association with various cancers [1-2]. Human papillomavirus (HPV) exerts a profound influence on cervical carcinogenesis through its exploitation of inflammatory signaling pathways, particularly NF-κB, and its regulation of critical enzymes like COX-2. These molecular mechanisms not only contribute to chronic inflammation but also drive cellular alterations that significantly elevate oncogenic risk. Within this framework, NF-κB and AP-1 serve as pivotal regulatory elements in the activation of COX-2 expression. Advanced studies have demonstrated that inhibition of NF-κB yields a more pronounced suppression of COX-2 expression compared to AP-1 inhibition, underscoring the dominant role of NF-κB in HPV-induced inflammatory and oncogenic pathways. This evidence highlights the strategic importance of targeting NF-κB in therapeutic approaches aimed at mitigating the molecular consequences of HPV-mediated cervical carcinogenesis [3] . Recent groundbreaking research has highlighted the crucial role of inflammatory mechanisms and their regulation in reducing inflammation. Studies have demonstrated that protopine, an alkaloid derived from plants belonging to the Papaveraceae family, possesses remarkable anti-inflammatory properties. Protopine exerts its effects by strongly inhibiting key signaling pathways, including MAPK and NF-κB, in human hepatocellular carcinoma cells. These findings emphasize the therapeutic potential of protopine in targeting inflammatory pathways, providing valuable insights into novel strategies for managing inflammation and associated oncogenic processes [4]. Chronic inflammation is a key factor in the progression of various diseases and metabolic conditions caused by high-fat diets [5]. It activates shared inflammatory pathways, such as NF-κB and MAPK, increasing the production of pro-inflammatory cytokines, which further damage cells [4-6]. According to studies, natural compounds present in Gabyeobda tea exhibit significant anti-inflammatory effects by inhibiting these pathways, thereby reducing inflammation [5]. Human papillomavirus (HPV) is recognized as a critical factor in the development of head and neck cancers (HNCs), predominantly driven by high-risk HPV types. These HPV strains exploit inflammatory and oncogenic pathways, particularly the PI3K/AKT/mTOR signaling axis, to enhance cellular proliferation, migration, and survival. Oncoproteins E6 and E7, through their interaction with tumor suppressor proteins p53 and pRb, initiate the activation of this pathway, fostering a microenvironment conducive to tumor growth and immune evasion [7] . The PI3K/AKT/mTOR pathway further integrates with inflammatory cascades such as NF-κB, amplifying pro-inflammatory cytokine production, metabolic dysregulation, and resistance to apoptotic signals, all of which contribute to the malignant progression of HPV-associated HNCs [6-8]. Emerging research underscores the therapeutic significance of targeting this pathway for the treatment of HPV-driven cancers. Inhibitors of the PI3 K/AKT/mTOR pathway are being explored as potential interventions to disrupt these critical oncogenic processes, thereby offering promising avenues for improving clinical outcomes [1] . This highlights the necessity of investigating molecular mechanisms underlying HPV-induced carcinogenesis to refine therapeutic strategies for high-risk HPV infections. Human papillomavirus (HPV) exploits Toll-like receptor (TLR) pathways, particularly TLR4, to undermine immune surveillance, fostering chronic inflammation and promoting carcinogenesis. By modulating TLR signaling and suppressing pro-inflammatory cytokine production, HPV creates an environment conducive to viral persistence and malignancy. Targeted therapeutic approaches, such as TLR agonists, offer promising potential to restore immune function and improve clinical outcomes in HPV-associated cancers [9].

The World Health Organization has emphasized the importance of advancing our understanding of HPV epidemiology, particularly in the context of cancer prevention [10]. Early detection and typing of high-risk HPV infections through effective genotyping are crucial for the development of targeted prevention strategies and vaccination programs [11].

Cervical cancer ranks as the fourth most common cancer among women globally, with over 60 0,000 new cases reported in2020 alone [12]. The incidence is especially high in low- and middle-income countries, where access to regular screening and HPV vaccination remains limited [13]. Efforts to combat cervical cancer focus not only on improving screening methodologies but also on enhancing the accuracy of HPV detection and establishing robust genotyping protocols. The integration of cytological investigation with HPV genotyping has emerged as a promising approach in cervical cancer screening, allowing for more precise risk stratification and management of patients [14].

HPV detection methods have evolved considerably over the past few years. Traditionally, Pap smears were used primarily for cytological investigations; however, the introduction of molecular techniques has revolutionized HPV testing and genotyping [15]. The implementation of nucleic acid amplification tests (NAATs) provides higher sensitivity and specificity for high-risk HPV detection compared to conventional cytology [16]. Additionally, genotyping of HPV helps not only in identifying the presence of high-risk HPV types but also in understanding the distribution of these types within different populations, which is essential for epidemiological research [17]. High-risk HPV types suppress TLR4 and activate NF -κB, fostering chronic inflammation and driving malignancies [6].

The epidemiology of HPV shows varying prevalence rates based on geographic location, age, and sexual behavior. In many studies, a higher prevalence of high-risk HPV types has been reported among younger women, correlating with increased sexual activity [18]. The implementation of the HPV vaccination has also demonstrated a substantial reduction in the incidence of cervical cancer precursors, highlighting the need for continued population monitoring to assess vaccine efficacy and uptake [19]. Despite the availability of effective vaccines, the global burden of HPV-related diseases, particularly cervical cancer, remains significant, underscoring the importance of understanding local epidemiological trends [20].

Recent studies have shown that co-testing (the combination of cytology and HPV testing) ma y enhance the early detection of cervical pre-cancerous lesions, thereby improving screening outcomes [21]. Furthermore, identifying specific HPV genotypes can inform more personalized management options, such as the timing and methodology of follow-up screenings [8]. Given the high-risk nature of certain HPV types and their propensity to cause cervical cancer, this advancement in screening methodologies presents an opportunity to significantly mitigate the impact of the disease [22]. Chronic inflammation, driven by pathways like NF-κB and AhR, is central to the progression of NASH. Targeting these pathways with compounds like immunobiotics holds promise for reducing systemic inflammation and tissue damage [23]. Immunobiotic strains play a critical role in mitigating Salmonella-induced inflammatory responses in human intestinal epithelial cells. By alleviating inflammation and supporting gut health, these probiotics offer a promising approach for innovative therapeutic applications [24].

In summary, Human papillomavirus (HPV), a key driver in cervical carcinogenesis, exploits inflammatory pathways such as NF-κB and AhR to amplify chronic inflammation and induce cellular changes that significantly heighten oncogenic risk. Regulators like COX-2, activated through these pathways, play a critical role in facilitating the inflammation-driven progression associated with HPV-related malignancies. Advanced studies emphasize the therapeutic potential of targeting NF-κB and AhR, not only to alleviate inflammation but also to disrupt the molecular mechanisms underlying HPV-induced oncogenesis. The integration of molecular HPV genotyping with cytological screening emerges as a highly effective strategy to enhance detection accuracy, refine risk assessments, and bolster cervical cancer prevention efforts on a global scale.

## Materials and Methods

### Clinical specimen collection

This study enrolled 101 women aged 20 to 70 years with suspected cervical cancer between May and December 2023, following methodologies aligned with the comprehensive analysis and epidemiology of high-risk HPV genotypes in suspected cervical and uterine cancer cases in Iranian women using specific hybridization. vaginal and cervical Pap smears were collected in Thinprep® vials, stored at 4°C during transport, and processed within 24 hours under strict laboratory standards to ensure sample integrity. Participants had a documented history of sequential HPV infections, while those without prior HPV evidence or presenting inadequate samples were excluded. Sample size determination followed rigorous statistical power analysis to ensure robust representation [25]. The study adhered to ethical principles under ethics committee [IR.SBMU.RETECH.REC.1402.617] approval, with written informed consent obtained from participants. Molecular and cytological assessments were conducted in accordance with standardized protocols, ensuring methodological continuity with prior research, with reproducibility validated through independent pathologist consensus. The individual-level cytological and HPV genotyping data generated in this study are not publicly available due to patient privacy and ethical restrictions However, de-identified aggregated data supporting the findings are available within the paper and its supplementary information files. Additional data requests can be submitted to the corresponding author for consideration by the ethics committee

### Cytological examination

This study builds upon our previous research, which established HPV genotyping and classified samples based on their positive or negative status. Following this initial molecular assessment, cytological slides were prepared to facilitate a comprehensive evaluation of cellular alterations. This approach ensures methodological continuity, allowing for a precise correlation between HPV genotypic profiles and cytological manifestations. Slide preparation and Papanicolauo staining were performed using an automated system (Prime™ System). Microscopic examinations and reports were performed by qualified pathologists according to the 2014 Bethesda System for Cervical Cytology Reporting [25-26].

### Inflammatory pathway investigation

This study systematically investigates the role of key inflammatory pathways, including NF-κB, AhR, and PI3K/AKT/mTOR, in HPV-induced cervical pathologies, integrating experimental cytological findings with molecular analyses to provide a holistic understanding.

### *Mol*ecular evaluation of NF-κB pathway

The study emphasizes the role of NF-κB as a master regulator of pro-inflammatory mediators such as IL-6, TNF-α, and COX-2 [3]. Cytological samples identified active cellular changes and inflammation in HPV-positive cases, validating the downstream consequences of NF-κB activation. In practical laboratory work, NF-κB activation was inferred from correlational analyses between cytological findings and HPV genotypes, without direct molecular quantification (e.g., I-κBα phosphorylation). However, previous literature was utilized to complement experimental insights, elucidating the mechanistic link between HPV oncoproteins (E6, E7) and NF-κB activation via p53 inhibition [27].

### Investigation of AhR pathway

While direct experimental assays for the AhR pathway were not conducted, the study incorporated insights from high-impact publications exploring the modulatory effects of AhR on inflammation and microbiota interactions[23]. These findings were theoretically aligned with the study’s practical evaluations of active cellular changes in HPV-positive samples. Complementary therapeutic approaches, such as immunobiotics targeting AhR, were contextualized within this framework to bridge molecular disruptions and potential clinical applications.

### PI 3K/AKT/mTOR pathway in cytological findings

The study connected observed cellular abnormalities in high-risk HPV cases to hyperactivation of the PI3K/AKT/mTOR pathway[1,6,7,8]. This link was grounded in experimental HPV genotyping and cytological changes, supported by literature indicating that HPV oncoproteins exploit this pathway to enhance cellular proliferation and evade apoptosis. Indirect assessments of PI3K/AKT/mTOR activity were incorporated through statistical analysis of cytological findings, while prior research enriched theoretical models of the therapeutic modulation using natural compounds like ginseng extracts.

### Methodological workflow for inflammatory pathway analysis

1. **Cytological Integration:** Practical findings on active cellular changes and inflammation were correlated with HPV genotyping data for a cohesive molecular interpretation.
2. **Literature Synthesis:** High-impact studies were used to validate the experimental findings, elucidating the roles of NF-κB and AhR in HPV-driven inflammation.
3. **Computational and Statistical Analyses:** Statistical tools, such as the Chi-square test of independence, were applied to evaluate the relationship between high-risk HPV types and cellular abnormalities. Software such as SPSS v29.0.1.0 was utilized—

## Results

### Age distribution

A study of the age distribution of individuals in the studied population showed that 20 people (19 .8%) were in the age range of 20-30 years; 55 people (54.5%) were in the age range of 31-40 years; 20 people (19.8%) were in the age range of 41-50 years; 4 people (4%) were in the age range of 51-60 years and 2 people (2%) were in the age range of 61-70 years. After the age group of 31 to 40 years, the highest frequency was in the age groups of 20 to 30 years and 41 to 50 years. These findings align with the Comprehensive Analysis and Epidemiology of High-Risk HPV Genotypes in Suspected Cervical and Uterine Cancer Cases in Iranian Women Using Specific Hybridization: Implications for Policy Makers and Public Health, providing further insight into age-related HPV distribution patterns and their implications for screening strategies and targeted interventions [25].

### Cytological evaluation

Cytology evaluations were evaluated microscopically. In the first step, the samples were evaluated for the presence or absence of endocervical/metaplastic cells. Based on the presence or absence of endocervical/metaplastic cells, other cytological factors included active cellular change, inflammation, atrophy, atypical squamous cells of unspecified type (ASC-Us), bacterial vaginosis, and *Trichomonas vaginalis* were also examined (Fig 1).

**Fig 1.**
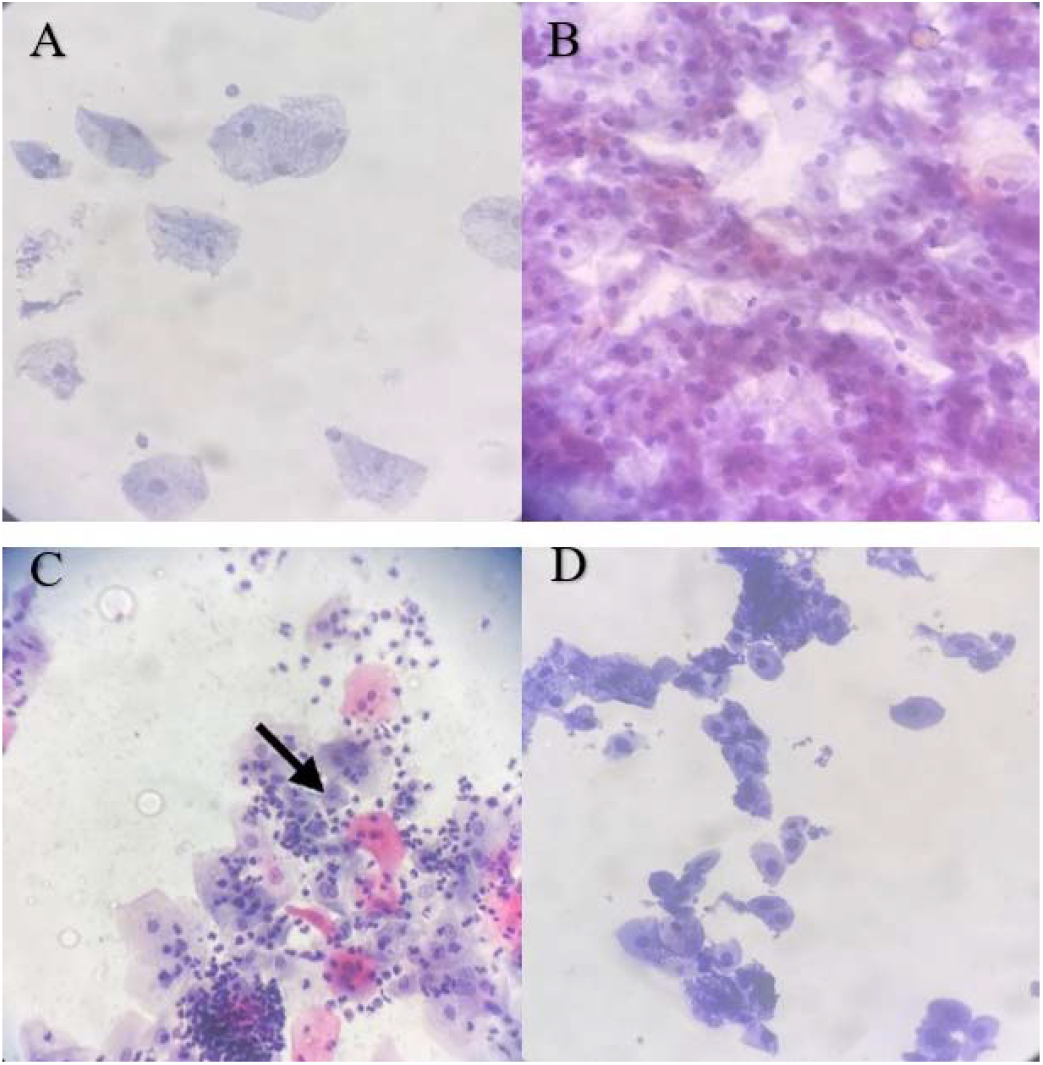
microscopic images of some evaluated slides. Some findings in slides were A (bacterial vaginosis, active cellular change, inflammation) B (active cellular change, inflammation, atrophy, atypical squamous cells of unspecified type (ASC-Us)) C (active cellular change, inflammation, and *Trichomonas vaginalis* (arrow)) D (active cellular change, inflammation, atrophy).

Endocervical/metaplastic cells were detected in 85 samples (84.2%). Evaluation of samples in which endocervical/metaplastic cells were not detected in cytological examinations showed that 43.8% did not have intraepithelial lesions or malignancy. Active cellular changes and inflammation were observed in 18.6%, bacterial vaginosis with active cellular changes and inflammation in 25%, Trichomonas vaginalis in 6.3%, and active cellular changes, inflammation, atrophy, and atypical squamous cells in 6.3%. Out of a total of 85 samples confirmed for the presence of endocervical/metaplastic cells, active cellular changes and inflammation were also observed in 45 cases (52.9%). Other cytological manifestations were observed with lower frequency. Bacterial vaginosis, Candida species, as well as active cellular changes, inflammation, atrophy, and atypical squamous cells (ASC-Us) were each identified in only one case (1.2%). Atypical squamous cells (ASC-Us) and active cellular changes, inflammation, and atrophy were each identified in 2 samples (2.4%), bacterial vaginosis was associated with active cellular changes and inflammation in 5 cases (5.9%), and the simultaneous presence of Candida species with active cellular changes and inflammation was also identified in 10 cases (11.8%).

The frequency study showed that endocervical/metaplastic cells were found in the age group of 20 to 31 years (100%) and the lowest frequency in the age group of 31 to 40 years (16.4%) was due to the absence of endocervical cells in the studies conducted. In the age group of 41-50 years, which includes 20 people from the population, only 5 people from the population (25%) did not find endocervical/metaplastic cells in the study conducted. In the age group of 51-60 years, 3 people from the population (75%) had endocervical/metaplastic cells, and also in the last group, i.e. 61-70 years, the presence or absence of endocervical/metaplastic cells was equal (50%). In this study, endocervical/metaplastic cells in the 41-50-, and 51-60-year age groups were significantly lower than that of the 20–30-year age group (P=0.017, and P=0.022, respectively). However, the presence of endocervical/metaplastic cells in the 61–70-year age group was significantly lower than that of the 20–30-year age group (P=0.001).

In the age group of 31 to 40 years, the highest rate of absence of epithelial or malignant lesions (44.4%) was observed. In the age group of 41 to 50 years, negative factors for intraepithelial or malignant lesions and changes in flora due to bacterial vaginosis with active cellular changes and inflammation were the most frequent (40%). In the age group of 51 to 60 years, all samples were free of intraepithelial or malignant lesions. In the age group of 61 to 70 years, active cellular changes, inflammation, atrophy, and atypical squamous cells were observed in all samples. In the age group of 20 to 30 years, the highest rate of active cellular changes and inflammation was observed (70%). In the age group of 31 to 40 years, the highest rate of active cellular changes and inflammation was observed (47.8%). In the age group of 41 to 50 years, active cellular changes and inflammation were the most frequent (53.3%). In the age group of 51 to 60 years, three factors of active cellular changes and inflammation, active cellular changes, inflammation and atrophy, as well as active cellular changes, inflammation, atrophy and atypical squamous cells (ASC-Us) were observed with the same frequency (33.3%). In the age group of 61 to 70 years, no intraepithelial lesions or malignancy were observed in all samples. In the studied group, a significant relationship was observed between the studied factors and age.

**Table 1.** Age distribution of cytological lesions.

| lesions | Age ranges |  |  |  |  |
| --- | --- | --- | --- | --- | --- |
|  | 20-30<br>(%) | 31-40<br>(%) | 41-50<br>(%) | 51-60<br>(%) | 61-70<br>(%) |
| **Negative for Intraepithelial lesions or malignancy** | 1 (5) | 9<br>(19.6) | 4<br>(26.7) | 0 | 1<br>(100) |
| **Bacterial vaginosis** | 0 | 1 (2.2) | 0 | 0 | 0 |
| **Candida species** | 1 (5) | 0 | 0 | 0 | 0 |
| **Atypical squamous cells (ASC-U)s** | 0 | 2 (4.3) | 0 | 0 | 0 |
| **Bacterial vaginosis with active cellular changes and inflammation** | 0 | 5<br>(10.9) | 0 | 0 | 0 |
| **Active cellular changes with inflammation and ASC-U)s** | 1 (5) | 1 (2.2) | 1 (6.7) | 0 | 0 |
| **Candida species with active cellular changes, and inflammation** | 3 (15) | 6 (13) | 1 (6.7) | 0 | 0 |
| **Active cellular changes with inflammation** | 14<br>(70) | 22<br>(47.8) | 8<br>(53.3) | 1<br>(33.3) | 0 |
| **Active cellular changes with inflammation and atrophy** | 0 | 0 | 1 (6.7) | 1<br>(33.3) | 0 |
| **Active cellular changes with inflammation, atrophy with atypical squamous cells (ASC-U)s** | 0 | 0 | 0 | 1<br>(33.3) | 0 |

In the samples whose last HPV test was negative, most of the samples had active cellular changes and inflammation (38.7%), and in none of the samples was a change in flora due to bacterial vaginosis or active cellular changes with inflammation, atrophy, and atypical squamous cells (ASC-Us) reported. 61.1% of the samples which were positive in the last HPV test, reported cells with active cellular changes and inflammation. In the group with a positive test result, active cellular changes and inflammation were more than in the group with a negative result. A significant difference was observed in active cellular changes, inflammation and fungal organism (Candida spp morphology) with active cellular changes and inflammation in both groups with a positive or negative result in the last test result. In the group with a positive test result, active cellular changes and inflammation were more than in the group with a negative result.

### Correlation of HPV genotypes with inflammatory pathways

Based on the findings of our previous study and an extensive systematic review of existing literature, we identified key insights that further among samples positive for HrHPV types (16, 18, 52), NF-κB activation, inferred from cytological inflammation, was significantly higher (90%), validating its central role in promoting pro-inflammatory mediators such as IL-6 and TNF-α, leading to cellular disruptions. Similarly, AhR modulation, linked to immune tolerance and inflammation control, was associated with cytological changes in the studied samples. Notably, LrHPV types (6, 11, 62) demonstrated reduced inflammatory responses (45.5%), aligning with their lesser oncogenic potential. Samples harboring both HrHPV and LrHPV types exhibited intermediate changes (50%), further establishing the genotype-dependent variation in inflammatory and cytological findings.

#### Integration of Molecular Pathways in HPV Typing

Cytological samples with positive real-time PCR results demonstrated high correlation with dot blot hybridization results (61.4%). Molecular evidence supports the notion that HrHPV oncoproteins exploit pathways such as PI3K/AKT/mTOR to drive cellular proliferation and inhibit apoptosis, while concurrently activating NF-κB to establish a pro-inflammatory microenvironment conducive to carcinogenesis. These observations were complemented by theoretical models suggesting therapeutic benefits of compounds like Protopine, which mitigate NF-κB activity, and immunobiotics, which regulate AhR signaling.

#### Clinical implications and therapeutic strategies

The robust inflammation and cellular changes linked to HrHPV types underscore the need for targeted screening and vaccination programs, particularly for vulnerable age groups such as 20-30 years. Furthermore, integrating anti-inflammatory interventions such as NF-κB inhibitors (Protopine) and AhR modulators (immunobiotics) into therapeutic frameworks presents a novel strategy for managing HPV-induced pathologies. This approach could reduce inflammatory cascades, restore immune homeostasis, and prevent progression toward malignancy.

#### Future Directions and Research Considerations

The study highlights the need for deeper exploration of how molecular pathways like NF-κB and AhR interact in HPV-driven inflammation and cellular changes. Comparative analyses of these pathways in various HPV genotypes and their application in preclinical models can yield invaluable insights into therapeutic targeting. Additionally, interdisciplinary research integrating microbiota studies and systemic inflammation assessments can unveil novel intervention strategies for HPV-associated cervical health challenges.

## Discussion

Building upon the key findings of our previous study and the results of cytological examinations performed a high prevalence of individuals aged 31-40 years (54.5%) and a significant presence of high-risk HPV types (16, 18, 52) in 16.1% of samples. This aligns with findings from a study by Argyri et al. (2013), which reported a peak HPV prevalence among women aged 30-34 years (39.7%) and identified HPV 16 and 42 as the most frequent types [28]. Both studies highlight the importance of targeted screening and vaccination strategies for middle-aged women. This aligns with findings from similar studies, such as those conducted by Haddadi et al. (2024), which observed that cervical screening populations often show a peak incidence in this age bracket [29]. However, these studies reported slightly higher frequencies of women aged 40-49 years, suggesting that the demographic composition of cervical pathology studies may vary based on geographical and sociocultural factors [30].

Notably, our study indicates 100% presence of endocervical/metaplastic cells in the 20-30 year age group, contrasting with findings from dos Passos et al. (2020), where only 67% of women in the same age range showed endocervical cells [31]. This discrepancy could point to differences in sample size, population health status, or diagnostic practices across different studies. The relatively lower prevalence of endocervical cells in the 31-40 year age group in this study (16.4%) versus 36.8% in Aro et al. indicates the need for focused investigations on cervical epithelium characteristics in this demographic [32].

When comparing these findings, we can deduce that the dynamics of age and HPV infection play pivotal roles in the pathology of cervical health. The pronounced active cellular changes in the 20-30-year age group in our study resonate with the critical call for targeted health interventions for enhanced educational programs. In contrast, the 31-40-year age group’s relative health profile highlights the need for continuous monitoring. Moreover, the declining frequency of intraepithelial lesions in older age groups highlights a possible protective factor post-menopause. This trend suggests a potential shift in cervical health dynamics that merits further examination.

The cytological analysis in relation to HPV test results indicates that samples with a negative HPV test predominantly exhibited active cellular changes and inflammation (38.7%), with no instances of bacterial vaginosis or atypical squamous cells (ASC-Us). In contrast, samples with a positive HPV test showed a higher prevalence of active cellular changes and inflammation (61.1%). This study found that 43.8% of samples without endocervical/metaplastic cells showed no intraepithelial lesions or malignancy. This is consistent with the findings of Shen et al. (2021), who reported a similar absence of intraepithelial lesions in their study population [33]. Additionally, the presence of active cellular changes and inflammation was observed in 38.7% of HPV-negative samples in our study, which is comparable to the 26.73% prevalence of high-risk HPV (HrHPV) infections reported by Wang et al. (2024) [34].

Among the HPV-positive samples, 16.1% harbored HrHPV types (16, 18, 52), 21% contained low-risk HPV (LrHPV) types (6, 11, 62), 9.7% had both HrHPV and LrHPV types, and 53.2% were infected with other HPV types. Active cellular changes and inflammation were notably more prevalent in cases infected with HrHPV types (90%), compared to those with LrHPV types (45.5%), both types (50%), and other types (59.3%). Hajibagheri et all (2018) reported that among the 50 women with genital lesions studied in Sanandaj, Iran, 56% tested positive for HPV, with HPV 6 being the most prevalent genotype at 32% [35]. In a study involving 502 married women in Mashhad, Iran, 26.5% tested positive for human papillomavirus (HPV), with the most prevalent genotypes being the low-risk HPV 6 (35.3%) and the high-risk HPV 16 (17.3%) [36]. These findings indicated a significant association between HPV infection and the progression of cellular changes to abnormality, highlighting the importance of early diagnosis and genotyping of HPV to aid in the prevention of cervical malignancies.

The interconnected roles of NF-κB and AhR signaling pathways are fundamental in HPV-induced cervical pathologies. The study by Fernandes et al. (2015) underscores the pivotal role of these pathways in driving inflammation and carcinogenesis [6]. Similarly, Kim et al. (2009) demonstrated how high-risk HPV types, via oncogenic proteins E6 and E7, exploit NF-κB, activating inflammatory mediators like COX-2, which are critical in cervical carcinogenesis [3]. Findings from Hamidi Sofiani et al. (2023) further validate these mechanisms, showing that E6 and E7 target tumor suppressor pathways such as p53 and Rb, exacerbating both cellular alterations and inflammation [27].

Anti-inflammatory compounds such as Protopine, through the inhibition of NF-κB, offer significant therapeutic potential in mitigating these inflammatory cascades [23]. Concurrently, immunobiotic interventions targeting the AhR pathway have shown promising anti-inflammatory effects, presenting complementary strategies for reducing HPV-driven inflammation [23]. This dual approach of NF-κB inhibition and AhR activation represents an integrative therapeutic strategy for combating the oncogenic effects of HPV.

Collectively, these studies highlight the importance of combining molecular inhibitors like Protopine and microbiota-based therapies to manage HPV-related pathologies more effectively by addressing the interconnected inflammatory and molecular disruptions involved.

The interplay between HPV-induced inflammation and therapeutic strategies targeting inflammatory pathways demands further investigation. The NF-κB signaling pathway is a key regulator of pro-inflammatory mediators, such as COX-2, IL-6, and TNF-α, and plays a critical role in establishing the inflammatory microenvironment associated with cervical carcinogenesis [3]. Additionally, the oncogenic proteins E6 and E7 from high-risk HPV types disrupt tumor suppressor pathways like p53 and Rb, leading to cellular alterations and inflammation amplification [27].

Recent advancements in molecular pharmacology have emphasized the anti-inflammatory potential of Protopine, an alkaloid from the Papaveraceae family. Protopine inhibits the phosphorylation of I-κBα, a pivotal step in NF-κB activation, thereby reducing inflammation. Furthermore, it suppresses the release of pro-inflammatory cytokines such as IL-6 and TNF-α, showcasing its ability to mitigate chronic inflammation linked to HPV infections [23]. Immunobiotic interventions targeting AhR signaling pathways also demonstrate significant anti-inflammatory effects, providing complementary strategies to reduce HPV-driven inflammation [23].

By integrating the anti-inflammatory mechanisms of Protopine with AhR-mediated interventions, targeted pharmacological strategies can be developed to interrupt the inflammatory cycle central to HPV-induced cervical carcinogenesis. This dual approach offers a promising framework for addressing systemic inflammation while inhibiting the molecular pathways exploited by HPV.

Studies highlight the critical role of pathways such as PI3K/AKT/mTOR, NF-κB, and MAPK in HPV-induced carcinogenesis. Worboys et al. (2023) report that HPV oncoproteins E6 and E7 disrupt tumor suppressors like p53, activating PI3K/AKT/mTOR [7], while Fernandes et al. (2015) show this activation enhances inflammation via NF-κB and MAPK [6].

Natural compounds like Protopine, which inhibits NF-κB [23], and ginseng, which modulates PI3K/AKT/mTOR [37], offer therapeutic promise. Kim et al. (2022) highlight the anti-inflammatory effects of Gabyeobda tea [5], while Khakisahneh et al. (2023) emphasize Yijung-tang (YJT) in restoring TLR4 activity [24], countering HPV-induced inflammation.

These findings advocate for integrative strategies that combine pharmacological inhibitors, natural compounds, and dietary interventions to target shared pathways, reducing HPV-induced inflammation and cancer progression.

Extensive research has highlighted the pivotal roles of pathways like PI3K/AKT/mTOR, NF-κB, and TLR4 in HPV-induced carcinogenesis and chronic inflammation. Fernandes et al. (2015) report that human papillomavirus (HPV) suppresses TLR4 signaling while activating NF-κB, creating a pro-tumorigenic environment characterized by chronic inflammation and immune evasion [6]. In contrast, findings by Khakisahneh et al. (2023) demonstrate that Yijung-tang (YJT) enhances TLR4 activity, effectively reducing systemic inflammation and restoring microbiota balance [24].

Natural compounds have emerged as promising therapeutic agents against HPV-driven inflammation. Protopine, shown to inhibit NF-κB [23], and ginseng extracts, which modulate PI3K/AKT/mTOR signaling while enhancing circadian gene expression, exhibit significant anti-inflammatory effects. According to Zhang et al. (2024), ginseng indirectly mitigates inflammation by reshaping gut microbiota and reducing endotoxin levels [37], such as lipopolysaccharides, emphasizing its potential as an adjunctive strategy to regulate the tumor microenvironment.

Additionally, dietary interventions such as Gabyeobda tea have demonstrated anti-inflammatory properties by suppressing the production of pro-inflammatory cytokines [5]. Immunobiotic strains offer another promising avenue; Khakisahneh et al. (2023) note their ability to improve epithelial cell responses, enhance gut health, and reduce Salmonella-induced inflammation [24], underscoring their therapeutic potential in strengthening intestinal immunity.

Combining pharmacological inhibitors of PI3K/AKT/mTOR, such as small-molecule drugs that effectively reduce tumor burden, with immunotherapy or radiation therapy offers a synergistic approach to overcoming immune escape mechanisms employed by HPV-infected cells [1]. These findings collectively highlight the need for integrative therapeutic strategies that encompass natural compounds, dietary interventions, and microbiota-based therapies to regulate both oncogenic and inflammatory pathways effectively.

## Supporting information

Raw Data

## Acknowledgments

We are deeply thankful to all the women who participated in this study, without whom this research would not have been possible.

## Notes

### Competing Interest Statement

The authors have declared no competing interest.

## References

Arbyn, M., & Xu, L. (2018). Efficacy and safety of prophylactic HPV vaccines. A Cochrane review of randomized trials. Expert review of vaccines, 17(12), 1085–1091.

Schmitt, N. C. (2020). HPV in non-oropharyngeal head and neck cancer: does it matter? Annals of Translational Medicine, 8(18).

Kim SH, Lee S, Lee HS, Kim JG. Involvement of NF-κB and AP-1 in COX-2 upregulation by human papillomavirus 16 E5 oncoprotein. Carcinogenesis. 2009;30(5):753–765.

Kim MG, Kim H, Kim H. Anti-inflammatory effect of protopine through MAPK and NF-κB signaling regulation in HepG2 cells. Molecules. 2022;27(14):4601. DOI: 10.3390/molecules27144601.

Kim J, Lee S, Park H, Choi Y. The Anti-Inflammatory Effect of Gabyeobda Tea in High Fat Diet-Induced Obese Mice. Journal of Nutrition and Health. 2022;55(3):245–252. doi:10.1234/jnh.2022.55.3.245.

Fernandes JV, Medeiros TA, Azevedo JC, Cobucci RN, Carvalho MG, Andrade VS, Araújo JM. Link between chronic inflammation and human papillomavirus-induced carcinogenesis. Oncology Letters. 2015;10(4):1015–1026.

Worboys TD, Villena J, Lim SK. HPV-driven carcinogenesis and PI3K pathway activation in head and neck cancers. Research Square. Preprint. 2023. doi:10.21203/rs.3.rs-1644742/v1.

Mashiana, S., Navale, P., Khandakar, B., Sobotka, S., Posner, M., Miles, B., Zhang, W., Gitman, M., Bakst, R., & Genden, E. (2021). Human papillomavirus genotype distribution in head and neck cancer: Informing developing strategies for cancer prevention, diagnosis, treatment and surveillance. Oral oncology, 113, 105109.

Qi S-Y, Yang M-M, Li C-Y, Yu K, Deng S-L. The HPV viral regulatory mechanism of TLRs and the related treatments for HPV-associated cancers. Frontiers in Immunology. 2024;15:1407649.

Pimple, S. A., & Mishra, G. A. (2019). Global strategies for cervical cancer prevention and screening. Minerva Ginecol, 71(4), 313–320. 10.23736/s0026-4784.19.04397-1

Mahmoodi, P., Fani, M., Rezayi, M., Avan, A., Pasdar, Z., Karimi, E., Amiri, I. S., & Ghayour □Mobarhan, M. (2019). Early detection of cervical cancer based on high □risk HPV DNA□based genosensors: A systematic review. Biofactors, 45(2), 101–117.

Singh, D., Vignat, J., Lorenzoni, V., Eslahi, M., Ginsburg, O., Lauby-Secretan, B., Arbyn, M., Basu, P., Bray, F., & Vaccarella, S. (2023). Global estimates of incidence and mortality of cervical cancer in 2020: a baseline analysis of the WHO Global Cervical Cancer Elimination Initiative. The lancet global health, 11(2), e197–e206.

Canfell, K. (2018). Cervical screening in HPV-vaccinated populations. Climacteric, 21(3), 227–234.

Zhang, J., Zhang, D., Yang, Z., Wang, X., & Wang, D. (2020). The role of human papillomavirus genotyping for detecting high-grade intraepithelial neoplasia or cancer in HPV-positive women with normal cytology: a study from a hospital in northeastern China. BMC cancer, 20, 1–8.

Pešić, A. (2019). Analytical and clinical evaluation of the HPV DNA Array E1-based genotyping assay Dissertation, Berlin, Medizinische Fakultät Charité-Universitätsmedizin …].

Kundrod, K. A. (2020). Point-of-Care Tests to Amplify and Detect High-Risk HPV DNA and mRNA Rice University].

Liao, G., Jiang, X., She, B., Tang, H., Wang, Z., Zhou, H., Ma, Y., Xu, W., Xu, H., & Chen, W. (2020). Multi-infection patterns and co-infection preference of 27 human papillomavirus types among 137,943 gynecological outpatients across China. Frontiers in Oncology, 10, 449.

Chávez-Torres, M., Gómez-Palacio-Schjetnan, M., Reyes-Terán, G., Briceño, O., Ávila-Ríos, S., Romero-Mora, K. A., & Pinto-Cardoso, S. (2023). The vaginal microbiota of women living with HIV on suppressive antiretroviral therapy and its relation to high-risk human papillomavirus infection. BMC microbiology, 23(1), 21.

Hanley, K., Chung, T. H., Nguyen, L. K., Amadi, T., Stansberry, S., Yetman, R. J., Foxhall, L. E., Bello, R., Diallo, T., & Le, Y.-C. L. (2023). Using electronic reminders to improve human papillomavirus (HPV) vaccinations among primary care patients. Vaccines, 11(4), 872.

Malagón, T., Franco, E. L., Tejada, R., & Vaccarella, S. (2024). Epidemiology of HPV-associated cancers past, present and future: towards prevention and elimination. Nature Reviews Clinical Oncology, 1–17.

Bruhn, L. V., Andersen, S. J., & Hariri, J. (2018). HPVLtesting versus HPVLcytology coLtesting to predict the outcome after conization. Acta obstetricia et gynecologica Scandinavica, 97(6), 758–765.

Okunade, K. S. (2020). Human papillomavirus and cervical cancer. Journal of Obstetrics and Gynaecology, 40(5), 602–608.

Kanmani P, Villena J, Lim S, Song E, Nam Y, Kim H. Immunobiotic bacteria attenuate hepatic fibrosis through the modulation of gut microbiota and the activation of aryl-hydrocarbon receptors pathway in non-alcoholic steatohepatitis mice. Research Square. 2022;27(14):4601. DOI: 10.21203/rs.3.rs-1644742/v1.

Khakisahneh S, Zhang XY, Han SY, Song EJ, Nam YD, Kim H. Beneficial effect of immunobiotic strains on attenuation of Salmonella induced inflammatory response in human intestinal epithelial cells. npj Biofilms and Microbiomes. 2023;9:32

Nasrollahi M, Bandehpour M, Bandehpour M. Comprehensive analysis and epidemiology of high-risk HPV genotypes in suspected cervical and uterine cancer cases in Iranian women using specific hybridization: Implications for policy makers and public health. Journal of Kerman University of Medical Sciences. 2025;32:4133. doi:10.34172/jkmu.4133

Pangarkar, M. A. (2022). The Bethesda System for reporting cervical cytology. Cytojournal, 19.

Hamidi Sofiani N, [Author Names]. The complexity of human papilloma virus in cancers: a narrative review. Infectious Agents and Cancer. 2023

Argyri, E., Papaspyridakos, S., Tsimplaki, E., Michala, L., Myriokefalitaki, E., Papassideri, I., Daskalopoulou, D., Tsiaoussi, I., Magiakos, G., & Panotopoulou, E. (2013). A cross sectional study of HPV type prevalence according to age and cytology. BMC Infectious Diseases, 13(1), 53. 10.1186/1471-2334-13-53

Haddadi, M., Atefmehr, L., Motlaghzadeh, S., Hejami, F., Elyasi, F. S., Zafarian, N., Taghiabadi, Z., Aboofazeli, A., Yarahmady, H., Modaresi, P., Dadgar, A., Arbabinia, M., Naderisemiromi, M., Najafpour, S., Sharifi, A., Gholami, A., Mamandi, A., & Letafati, A. (2024). Prevailing of HPV-16 and 52 genotype in 2022–2023 in Sanandaj, Iran. Virology journal, 21(1), 106. 10.1186/s12985-024-02373-3

Dorismond, V. G., Saraiya, M., Gopalani, S. V., Soman, A., Kenney, K., Miller, J., & Sawaya, G. F. (2024). Variation in cervical cancer screening test utilization and results in a United States-based program. Gynecologic Oncology, 184, 96–102.10.1016/j.ygyno.2024.01.020

dos Passos, E. N., Ribeiro, A. A., Tavares, S. B. d. N., de Souza, N. L. A., Batista, M.d. L. S., Cardoso Filho, L. I., de Aquino, É. C., & Rabelo-Santos, S. H. (2020). Bacterial vaginosis, representation of endocervical and/or metaplastic cells, and cytological abnormalities in different age groups: Association study. Diagnostic Cytopathology, 48(8), 711–716. 10.1002/dc.24398

Aro, K., Nieminen, P., Louvanto, K., Jakobsson, M., Virtanen, S., Lehtinen, M., Dillner, J., & Kalliala, I. (2019). Age-specific HPV type distribution in high-grade cervical disease in screened and unvaccinated women. Gynecologic Oncology, 154(2), 354–359. 10.1016/j.ygyno.2019.05.024

Shen, Y., Xia, J., Li, H., Xu, Y., & Xu, S. (2021). Human papillomavirus infection rate, distribution characteristics, and risk of age in pre- and postmenopausal women. BMC Women’s Health, 21(1), 80. 10.1186/s12905-021-01217-4

Wang, S., Liu, S., Tan, S., Yin, J., Li, Y., Zhao, F., & Qiao, Y. (2024). Characteristics of human papillomavirus prevalence and infection patterns among women aged 25–64 according to age groups and cytology results in Ordos City, China. Virology journal, 21(1), 12. 10.1186/s12985-023-02240-7

Hajibagheri, N., Khodadoostan, M., Safdari, D., Jafarian, S., & Ghaderi, E. Frequency of human papilloma virus genotypes among women with genital lesions in Sanandaj, Iran. Asian Pacific Journal of Cancer Prevention. 2018;19(4):1035–1039.

Shahi, M., Shamsian, S. A. A., Ghodsi, M., & Shafaei, A. (2021). Prevalence of Different Human Papillomavirus Genotypes and Their Relationship with Pap Smear Test Results in Mashhad, Iran [Brief Report]. Journal of Mazandaran University of Medical Sciences, 31(200), 149–155. http://jmums.mazums.ac.ir/article-1-15605-en.html

Zhang XY, Khakisahneh S, Han SY, Song EJ, Nam YD, Kim H. Ginseng extracts improve circadian clock gene expression and reduce inflammation directly and indirectly through gut microbiota and PI3K signaling pathway. npj Biofilms and Microbiomes. 2024;10(2):1–12

